# Bioactivity-driven discovery of repurposable antivirals as OSCAR inhibitors that promote cartilage protection via transcriptomic reprogramming

**DOI:** 10.64898/2026.02.24.707642

**Authors:** Gina Ryu, Jihee Kim, Sera Park, Soo Young Lee, Wankyu Kim

**Affiliations:** Department of Life Science, Ewha Womans University, Seoul, Republic of Korea; The Research Center for Cellular Homeostasis, Ewha Womans University, Seoul, Republic of Korea; KaiPharm, Seoul 03760, Republic of Korea; Multitasking Macrophage Research Center, Ewha Womans University, Seoul, Republic of Korea

**Author notes:** These authors contributed equally: Gina Ryu, Jihee Kim, Sera Park.

## Abstract

Osteoarthritis (OA) is a progressive degenerative joint disorder characterized by cartilage degradation, chronic pain, and impaired joint function. The avascular nature of cartilage isolates chondrocytes from systemic circulation, presenting significant challenges for therapeutic intervention. Despite extensive efforts, no clinically effective disease-modifying osteoarthritis drugs (DMOADs) are currently available. Targeting chondrocyte-specific receptors has therefore emerged as a promising strategy. The osteoclast-associated receptor (OSCAR), expressed on chondrocytes, has been implicated in the regulation of cartilage homeostasis and OA pathogenesis.

Here, we applied sBEAR (Structurally similar Bioactive compound Enrichment by Assay Repositioning), a bioactivity-driven virtual screening framework independent of target structural information, to identify small-molecule inhibitors of the OSCAR–collagen interaction. By mining large-scale bioactivity profiles, we identified adefovir (ADV) and brivudine (BRV), as candidate OSCAR inhibitors. Molecular docking analyses indicated that both compounds occupy the collagen-recognition pocket within the OSCAR D2 domain.

Intra-articular administration of these compounds in a post-traumatic OA mouse model significantly attenuated OA progression and enhanced chondrocyte regeneration. Both compounds increased *Sox9* expression, and transcriptomic analyses revealed that BRV reverses inflammatory and extracellular matrix–degrading transcriptional programs.

Together, these findings establish OSCAR as a therapeutically actionable target in OA and highlight ADV and BRV as potential DMOAD candidates.

## Introduction

Osteoarthritis (OA) is a chronic and progressive degenerative joint disorder characterized by the degradation of articular cartilage, osteophyte formation, subchondral bone remodeling, and synovial inflammation^1^. As the most prevalent degenerative joint disease globally, OA represents a significant public health burden, being the primary cause of disability and reduced quality of life among aging populations^2^. Despite its widespread prevalence and profound socioeconomic impact, current therapeutic interventions are largely palliative, focusing on pain management rather than addressing the underlying pathophysiology^3^. This highlights an urgent need for the development of disease-modifying osteoarthritis drugs (DMOADs) capable of decelerating disease progression and promoting the restoration of joint structure and functionality^4^.

A defining feature of OA is the progressive degradation of the cartilage extracellular matrix (ECM)^5^. Under normal physiological conditions, chondrocytes sustain a delicate equilibrium between ECM synthesis and degradation through the production of both anabolic and catabolic mediators^6^. In OA, however, this homeostasis is disrupted, resulting in a pathological shift toward ECM catabolism^7^ driven by the upregulation of matrix metalloproteinases (MMP3, MMP9, and MMP13) and the aggrecanase ADAMTS5, which enzymatically degrade critical structural components of cartilage, including type II collagen and aggrecan (ACAN)^8^. Simultaneously, the biosynthesis of these essential ECM macromolecules is markedly diminished^9^. As such, targeting the molecular mechanisms that underpin this catabolic dysregulation represents a promising therapeutic avenue for mitigating OA progression^10^.

The osteoclast-associated receptor (OSCAR) is a receptor known to bind the triple-helical peptide sequence GPOGPAGFO within type II collagen through its extracellular domain^11^. While predominantly expressed in osteoclasts, where its interaction with collagen-II activates signaling cascades that co-stimulate FcRγ and facilitate osteoclastogenesis, its expression in normal chondrocytes is typically minimal^12^. However, studies have revealed that OSCAR expression is significantly upregulated in osteoarthritic chondrocytes in both human and murine models^13^. Furthermore, recent investigations have elucidated a direct role for OSCAR in OA pathogenesis: overexpression of OSCAR in murine joints via adenoviral vector delivery (Ad-OSCAR) was sufficient to recapitulate OA-like phenotypes, underscoring its critical involvement in disease progression^14^.

Based on this evidence, we postulated that small-molecule inhibitors capable of disrupting OSCAR’s interaction with type II collagen could represent promising candidates for DMOADs. However, targeting OSCAR with small molecules remains challenging, as its interaction with collagen is mediated by a relatively extended surface interface rather than a deep, well-defined binding pocket^15^, complicating conventional structure-based docking approaches. Moreover, the scarcity of well-characterized small-molecule ligands further limits traditional ligand-based discovery strategies.

To overcome these challenges, we employed a data-driven virtual screening strategy that leverages large-scale bioactivity datasets. Building on our previously developed BEAR (Bioactive compound Enrichment by Assay Repositioning) approach^16^, we further advanced sBEAR (Structurally Similar BEAR), which integrates chemical similarity–based enrichment to prioritize candidate compounds even in the absence of direct assay annotations. This approach enables efficient hit prioritization for targets with limited structural or bioactivity information.

Applying sBEAR to OSCAR, we systematically prioritized small molecules predicted to interfere with OSCAR–collagen interactions. Among the prioritized candidates, adefovir (ADV) and brivudine (BRV) emerged as representative compounds with favorable bioactivity profiles. Both ADV and BRV are clinically approved antiviral agents—ADV is used in the treatment of chronic hepatitis B^17^, while BRV is prescribed for herpes zoster^18^—highlighting their potential for drug repositioning. To support the feasibility of small-molecule modulation of OSCAR, we evaluated prioritized candidate compounds using molecular docking simulations. These analyses suggested that structurally diverse ligands can engage conserved interaction features within the OSCAR D2 domain associated with collagen recognition, consistent with a potential competitive mode of inhibition.

Guided by these computational insights, we next investigated whether pharmacological targeting of OSCAR could modulate disease-relevant phenotypes in osteoarthritis. We show that intra-articular (IA) administration of these compounds is associated with ameliorated disease outcomes–in a surgically induced OA mouse model, even when treatment is initiated at an advanced stage of disease. We further assessed the molecular consequences of OSCAR inhibition using integrated in vitro assays and transcriptomic profiling focused on a representative lead compound. These analyses revealed coordinated modulation of disease-relevant programs, including suppression of catabolic and inflammatory pathways and activation of chondrogenic differentiation and ECM restoration.

Collectively, these findings underscore the pivotal role of OSCAR in OA pathogenesis and highlight the potential of ADV and BRV as promising intra-articularly administered disease-modifying osteoarthritis drugs (DMOADs).

## Results

### Large-scale bioactivity–guided virtual screening identifies OSCAR-inhibitory candidate compounds

To identify candidate small-molecule inhibitors of OSCAR, we applied sBEAR, a data-driven virtual screening platform that leverages large-scale bioactivity profiles (Figure 1a). The sBEAR framework covered a screening space of 1,231,461 compounds, enabling bioactivity-based prioritization for challenging targets such as OSCAR.

**Figure 1.**
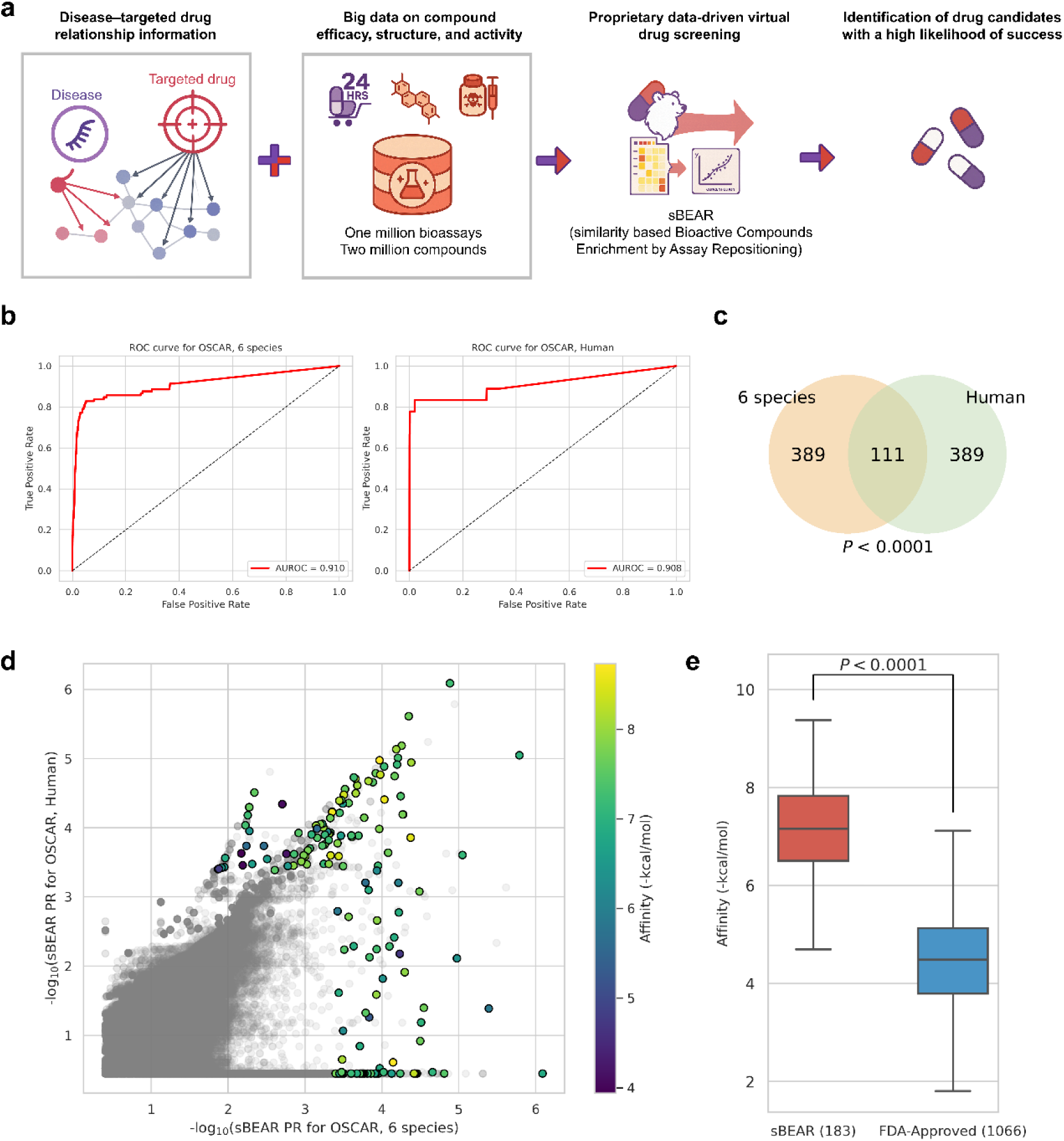
sBEAR-based virtual screening of candidate OSCAR inhibitors. (a) Schematic overview of the sBEAR framework used to prioritize compounds predicted to interfere with the OSCAR-collagen interaction based on large-scale bioactivity profiles. (b) Receiver operating characteristic (ROC) curves showing predictive performance of sBEAR models trained on known OSCAR ligands from six mammalian species (left) and human-specific ligands (right). (c) Venn diagram illustrating the overlap of the top 500 ranked predicted ligands between the six mammalian species and humans. (d) Percentile rank comparison of 183 curated candidate ligands between multi-species (x-axis) and human-specific (y-axis) predictions, with points colored according to predicted binding affinity. (e) Comparison of predicted binding affinities between 183 curated candidate compounds and a reference set of FDA-approved drugs. Statistical significance was assessed using the Mann–Whitney U test.

To evaluate screening performance, we assessed known OSCAR ligands using sBEAR. In 3-fold cross-validation, sBEAR demonstrated robust predictive performance (AUC = 0.908–0.910; Figure 1b) across OSCAR ligand sets derived from human and six mammalian species. The top 500 ranked candidates from each ligand set exhibited a significant overlap of 111 common candidates (Figure 1c). After removing redundant entries, including salt forms, stereoisoforms, and common metabolites, 183 unique candidates remained for further validation (Supplementary Table 1).

Ranking consistency between the multi-species and human models was further supported by comparison of percentile ranks, which showed concordant prioritization of curated candidate ligands, with many highly ranked compounds also exhibiting favorable predicted binding affinities (Figure 1d).

In addition, binding affinity prediction^19^ indicated that the curated candidate set exhibited significantly stronger predicted binding affinities to OSCAR compared with a reference set of FDA-approved drugs (Figure 1e), supporting effective enrichment of OSCAR-interacting compounds by the sBEAR pipeline.

### Experimental validation of ADV and BRV as dual-function OSCAR inhibitors

Subsequently, we moved on to experimental validation of OSCAR binding activity for the readily purchasable 45 compounds, which subjected to ELISA competition assay to assess binding affinity (Figure 2a, Supplementary Table 2). The ELISA assay evaluates the ability to inhibit the interaction between OSCAR and collagen, a known ligand of the OSCAR receptor^11^. Based on the degree of binding inhibition, secondary target validation narrowed the pool to 12 top-ranking candidate molecules. Subsequently, the top 12 candidate compounds were evaluated in chondrocytes stimulated with the pro-inflammatory cytokine IL-1β. Following IL-1β treatment, each compound was administered to assess its ability to suppress the IL-1β–induced upregulation of MMP3 mRNA expression. Based on the degree of MMP3 inhibition, five compounds showing the most potent effects were identified as second-hit candidates (Figure 2b).

**Figure 2.**
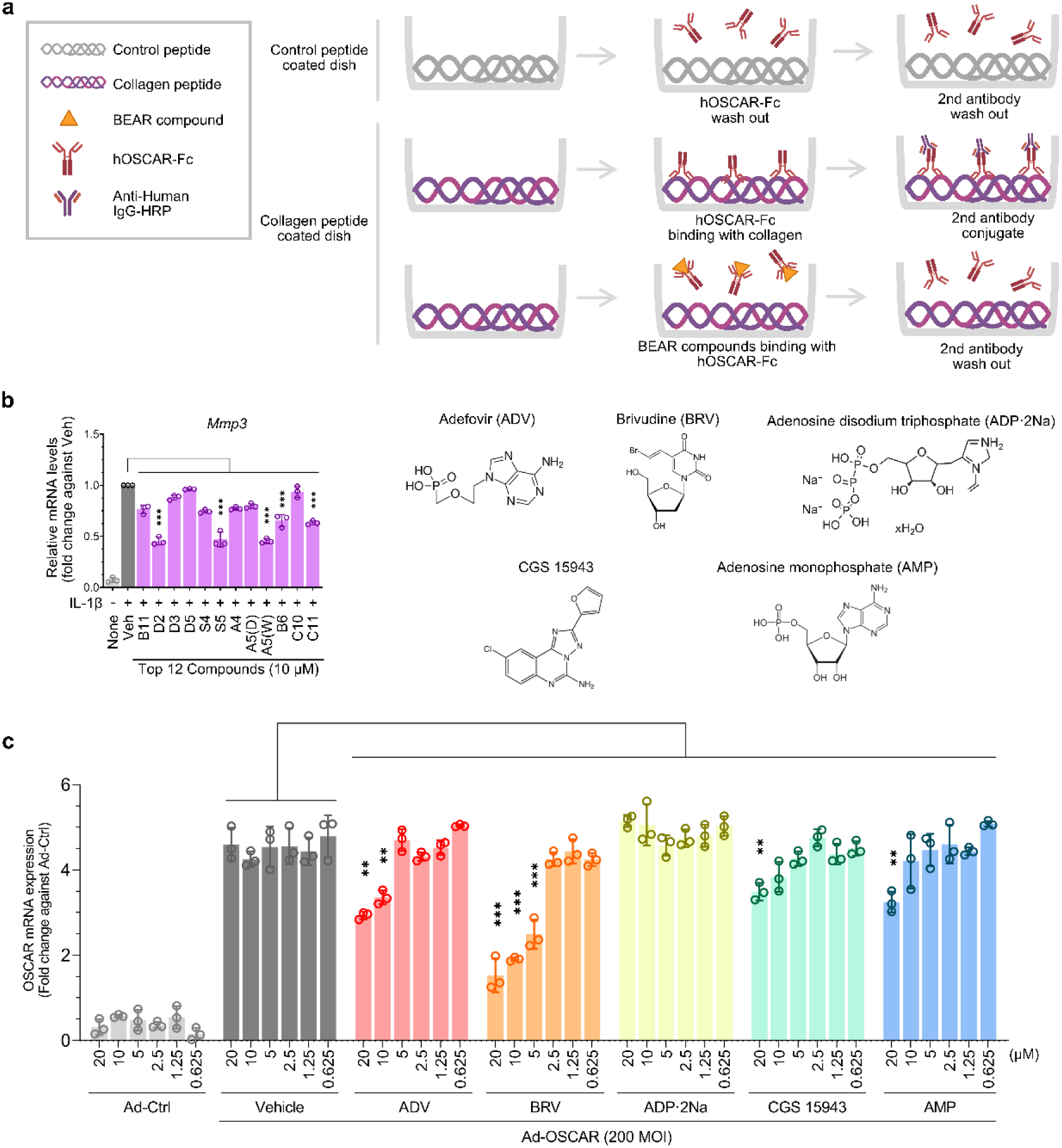
Stepwise screening strategy for identifying dual-function OSCAR inhibitors. (a) ELISA-based secondary screening narrowed down the initial 45 hit compounds to 12 candidates by measuring inhibition of OSCAR–collagen binding. (b) The top 12 compounds were further evaluated in IL-1β–stimulated chondrocytes, and five compounds—Adefovir (ADV), Brivudine (BRV), Adenosine disodium triphosphate (ADP·2Na), CGS 15943, and Adenosine monophosphate (AMP)—were selected based on their potent suppression of MMP3 mRNA expression. (c) Adenoviral OSCAR overexpression assay in chondrocytes was performed to assess the ability of the five selected compounds to suppress OSCAR expression; two compounds, Adefovir (ADV) and Brivudine (BRV), significantly reduced OSCAR levels.

To further validate the functional relevance of the top candidates, we performed an Adeno-OSCAR overexpression assay in chondrocytes to identify compounds capable of not only inhibiting OSCAR–collagen binding but also suppressing OSCAR receptor overexpression. Among the five top-ranking candidates, ADV and BRV significantly reduced OSCAR expression levels. These two compounds demonstrated multiple effects in chondrocytes, including inhibition of OSCAR–collagen interaction, suppression of IL-1β–induced MMP3 expression, and downregulation of OSCAR overexpression. Based on these multifaceted activities, ADV and BRV were selected as the final lead compounds (Figure 2c).

Together, these results highlight a stepwise screening strategy that successfully identified ADV and BRV as dual-function inhibitors of OSCAR, capable of modulating both ligand binding and receptor expression in chondrocytes, thereby positioning them as promising therapeutic candidates for cartilage-protective interventions.

### Structural interpretation of ADV and BRV engagement at the OSCAR collagen-recognition surface

For structural interpretation on how candidate compounds may interfere with OSCAR–collagen recognition, we performed structural docking simulations for the top-ranked compounds^20^. Docking analyses focused on ADV and BRV, which emerged as final lead candidates based on the experimental outcomes.

Both ADV and BRV were consistently predicted to bind within a surface-exposed groove located in the OSCAR D2 domain (Figure 3a), corresponding to the primary collagen-recognition site with relatively shallow surface enriched in polar and aromatic residues, rather than a well-defined binding pocket^15^. Structural studies have shown that, unlike related collagen receptors such as GPVI and LAIR-1, OSCAR engages collagen predominantly through its D2 domain, where an extended groove accommodates the triple-helical collagen motif. The predicted binding poses of ADV and BRV spatially overlapped with this D2 groove, suggesting a plausible mechanism by which these compounds may interfere with collagen engagement.

**Figure 3.**
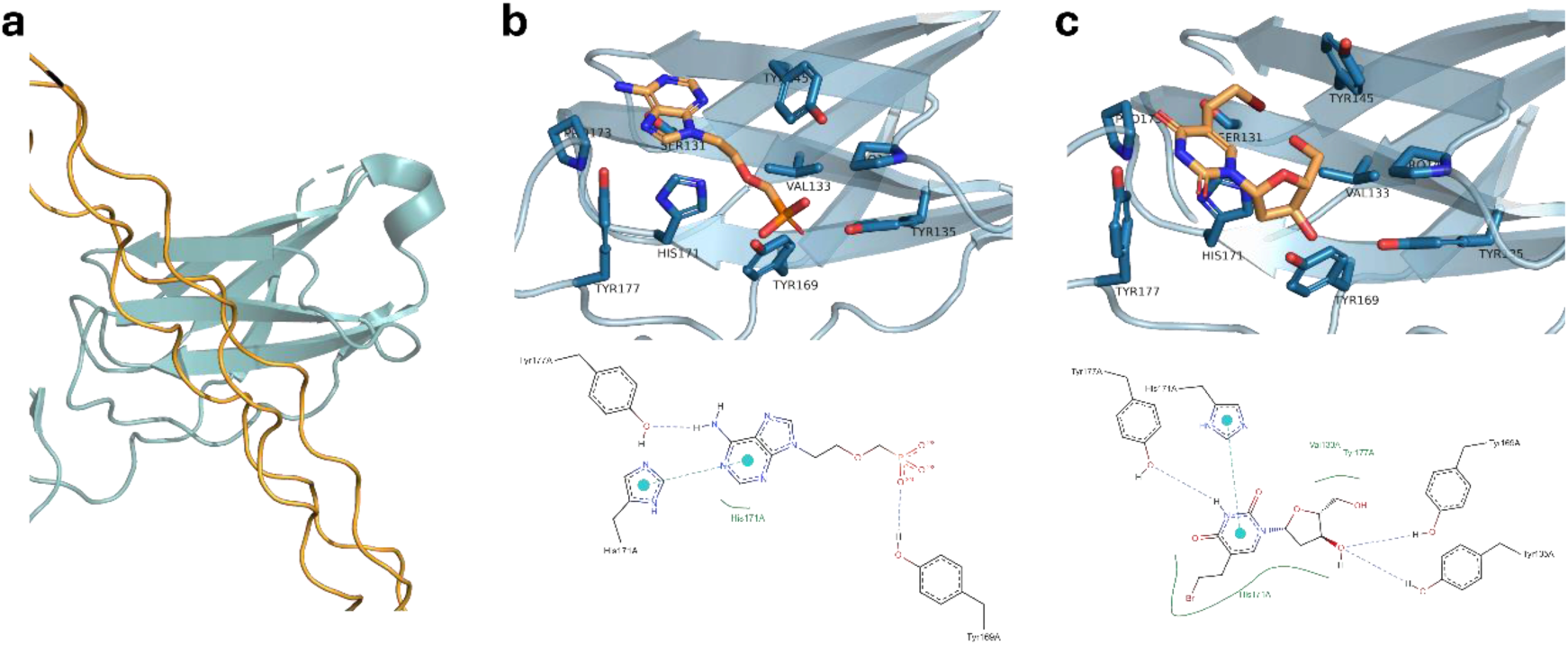
Structural context of small-molecule engagement at the OSCAR collagen-recognition surface. (a) Crystal structure of the OSCAR D2 domain in complex with a collagen triple-helical peptide, highlighting the surface that mediates collagen recognition. Predicted binding poses (top) and corresponding two-dimensional interaction maps (bottom) of (b) BRV, (c) ADV within the collagen-recognition surface of the OSCAR D2 domain. The docking poses overlap with the region implicated in collagen binding, suggesting a potential competitive mode of interference.

Despite their distinct chemical scaffolds, ADV and BRV displayed convergent interaction patterns within the D2 groove. Several residues previously implicated in collagen binding—corresponding to conserved aromatic and polar residues lining the D2 surface—were recurrently involved in ligand interactions. ADV formed hydrogen bonds with residues corresponding to His171 and Tyr177, along with electrostatic interactions involving the phosphate moiety, consistent with stabilization within a polar microenvironment (Figure 3b). Additional aromatic contacts with tyrosine residues further contributed to ligand positioning.

BRV similarly engaged multiple aromatic residues within the D2 groove through hydrogen bonding and π–π stacking interactions (Figure 3c). The presence of a brominated nucleobase enabled additional polarizable interactions, consistent with accommodation within the chemically versatile binding surface described for the OSCAR D2 domain.

These interaction patterns are consistent with the observation that the OSCAR D2 domain engages collagen via an extended surface enriched in polar and aromatic residues, rather than through a deep, enclosed pocket. Consistent with these observations, docking simulations of additional top-ranked candidates revealed recurrent localization to the OSCAR D2 collagen-recognition surface (Supplementary Figure 1), supporting the generality of this binding mode.

Collectively, these results support the feasibility of small-molecule engagement of the OSCAR collagen-recognition surface and are consistent with prior structural evidence defining D2 as the dominant functional interface for ligand recognition. Together, these predicted spatial relationships provide a structural framework that aligns with the observed inhibition of OSCAR–collagen interaction in biochemical and cellular assays.

### Protective effects of ADV and BRV on chondrocyte homeostasis under inflammatory conditions

Building upon our previous findings that modulation of OSCAR expression in chondrocytes can influence chondrocyte fate^14^, we further investigated whether treatment with ADV and BRV could regulate this process (Figure 4a). Chondrocytes were treated with the compounds, and their effects on chondrocyte fate were assessed.

**Figure 4.**
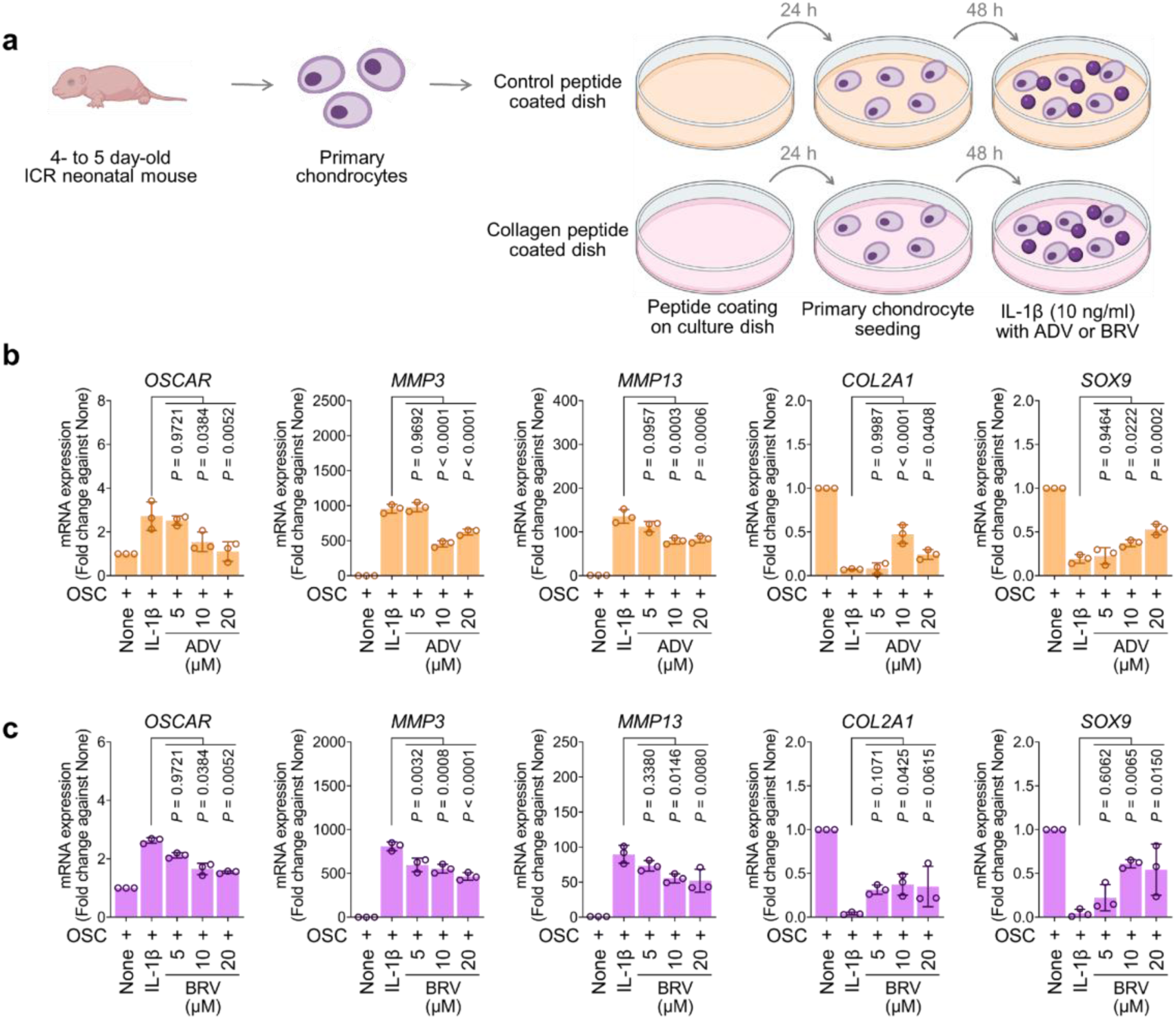
ADV and BRV protect chondrocytes from IL-1β–induced catabolic stress by modulating both catabolic and anabolic gene expression. (a) Schematic overview of the experimental approach used to evaluate the effects of Adefovir and Brivudine on chondrocyte fate under inflammatory conditions. (b) In IL-1β–stimulated chondrocytes, treatment with ADV significantly reduced the expression of catabolic markers (OSCAR, MMP3, MMP13) and restored the expression of anabolic markers (Col2a1, Sox9). (c) Similarly, BRV treatment suppressed catabolic gene expression and promoted anabolic marker recovery in chondrocytes exposed to IL-1β.

In chondrocytes stimulated with IL-1β, a pro-inflammatory cytokine known to induce the expression of matrix-degrading enzymes, treatment with ADV significantly reduced the expression of catabolic markers (e.g., OSCAR, MMP3, MMP13) and restored the expression of anabolic markers (e.g., Col2a1, Sox9), as shown in Figure 4b. Similarly, treatment with BRV also suppressed catabolic gene expression while promoting anabolic marker recovery in IL-1β–stimulated chondrocytes (Figure 4c). These results suggest that ADV and BRV not only inhibit OSCAR receptor function but also exert protective effects on chondrocytes by attenuating the catabolic response and promoting anabolic activity in an inflammatory environment.

### ADV and BRV Attenuate OA Progression by Preserving Cartilage and Subchondral Bone Integrity

To assess whether ADV and BRV could alleviate DMM-induced osteoarthritis (OA), we administered intra-articular injections of either 0.5 or 2 mg/kg of each compound for 8 weeks, beginning one week after surgery. Mice were sacrificed at week 9 for analysis (Figure 5a).

**Figure 5.**
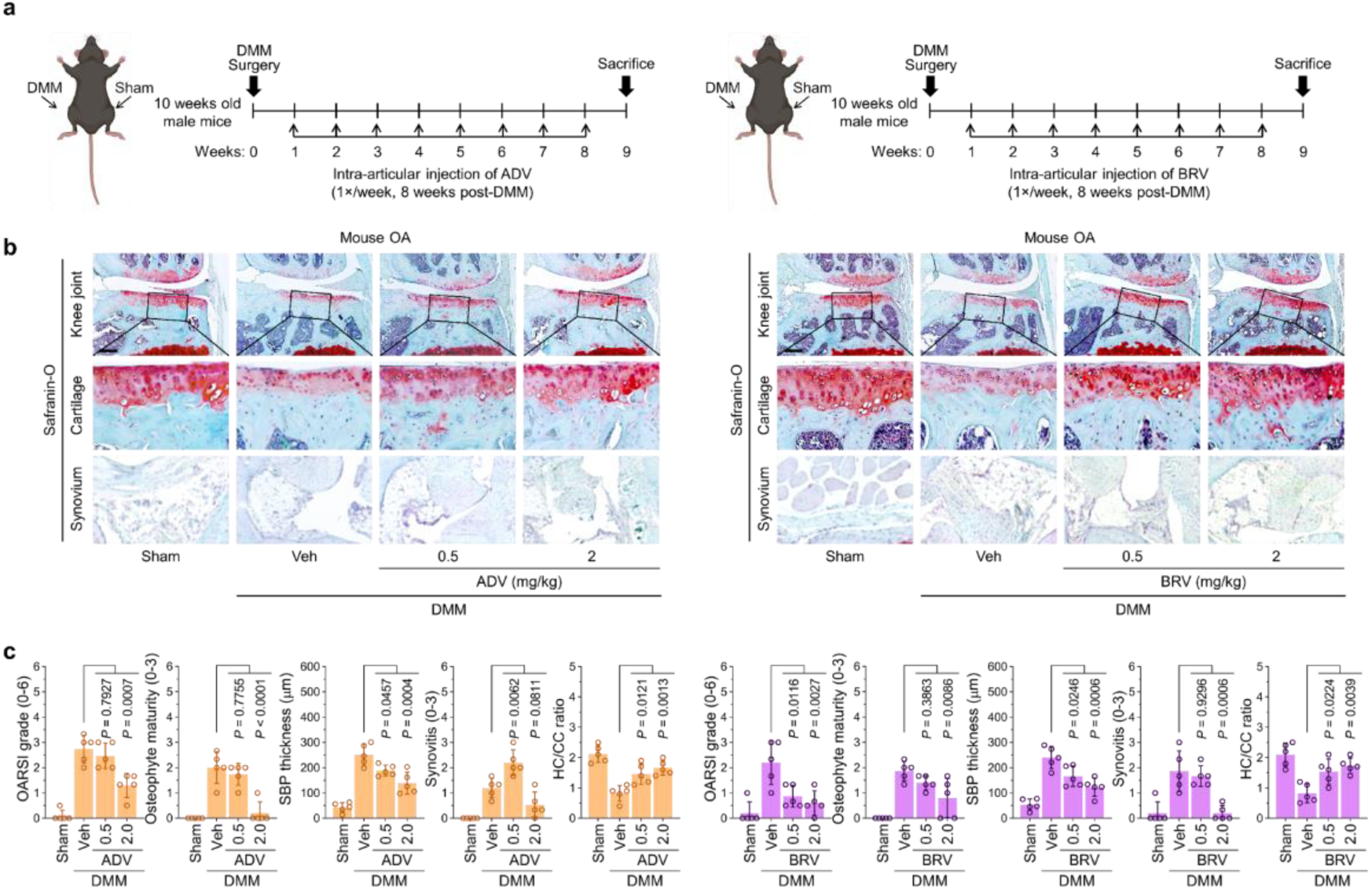
Intra-Articular ADV and BRV Protect Against DMM-Induced Osteoarthritis in a Dose-Dependent Manner. (a) Experimental timeline: mice underwent DMM surgery, followed by intra-articular injections of ADV or BRV (0.5 or 2 mg/kg) once weekly for 8 weeks, beginning 1 week post-surgery. Mice were sacrificed at week 9 for analysis. (b) Representative histological images and quantification showing dose-dependent reduction in articular cartilage erosion following treatment with ADV or BRV. (c) Quantitative analysis demonstrating that 2 mg/kg of adefovir or brivudine significantly improved cartilage integrity (OARSI score), osteoblast maturation, subchondral bone plate (SBP) thickness, and synovial inflammation. The OARSI grade, synovitis and osteophyte maturity data are shown as means ± 95% confidence intervals (CI). Differences between groups were determined with Kruskal-Wallis test followed by Mann–Whitney *U* test. Means ± s.e.m. with two-tailed *t*-test for SBP thickness. Exact *P* values can be found in the accompanying Source Data. Scale bars, 25 μm.

Both ADV and BRV dose-dependently reduced articular cartilage erosion in the knee joint (Figure 5b). Notably, treatment with 2 mg/kg of either compound significantly improved postoperative cartilage integrity, as evidenced by lower OARSI scores. In addition, this dosage reduced osteophyte maturation, decreased subchondral bone plate (SBP) thickness, and reduced synovial inflammation (Figure 5c). These findings suggest that intra-articular administration of ADV and BRV effectively protects against OA progression by preserving cartilage integrity and improving subchondral bone and synovial health.

### BRV Enhances Chondrogenic Differentiation of MSCs and Supports Cartilage Regeneration in Osteoarthritic Conditions

Based on our previous findings that intra-articular administration of ADV and BRV effectively suppressed the progression of moderately advanced osteoarthritis (OA) following DMM surgery, we further investigated the potential of these compounds to promote cartilage regeneration. Specifically, we examined whether ADV and BRV could induce the chondrogenic differentiation of rat bone marrow-derived mesenchymal stem cells (MSCs) into glycosaminoglycan (GAG)–producing chondrocytes (Figure 6a). Notably, both BRV and ADV robustly and dose-dependently enhanced chondrogenic differentiation, as evidence by increased Alcian Blue staining (Figure 6b, 6c). IC₅₀ analysis of the catabolic marker MMP3 revealed values of 0.48 μM for ADV and 7.5 μM for BRV (Figure 6d). These findings suggest that BRV, in particular, holds therapeutic potential for supporting cartilage regeneration in osteoarthritic joints by promoting chondrogenic differentiation.

**Figure 6.**
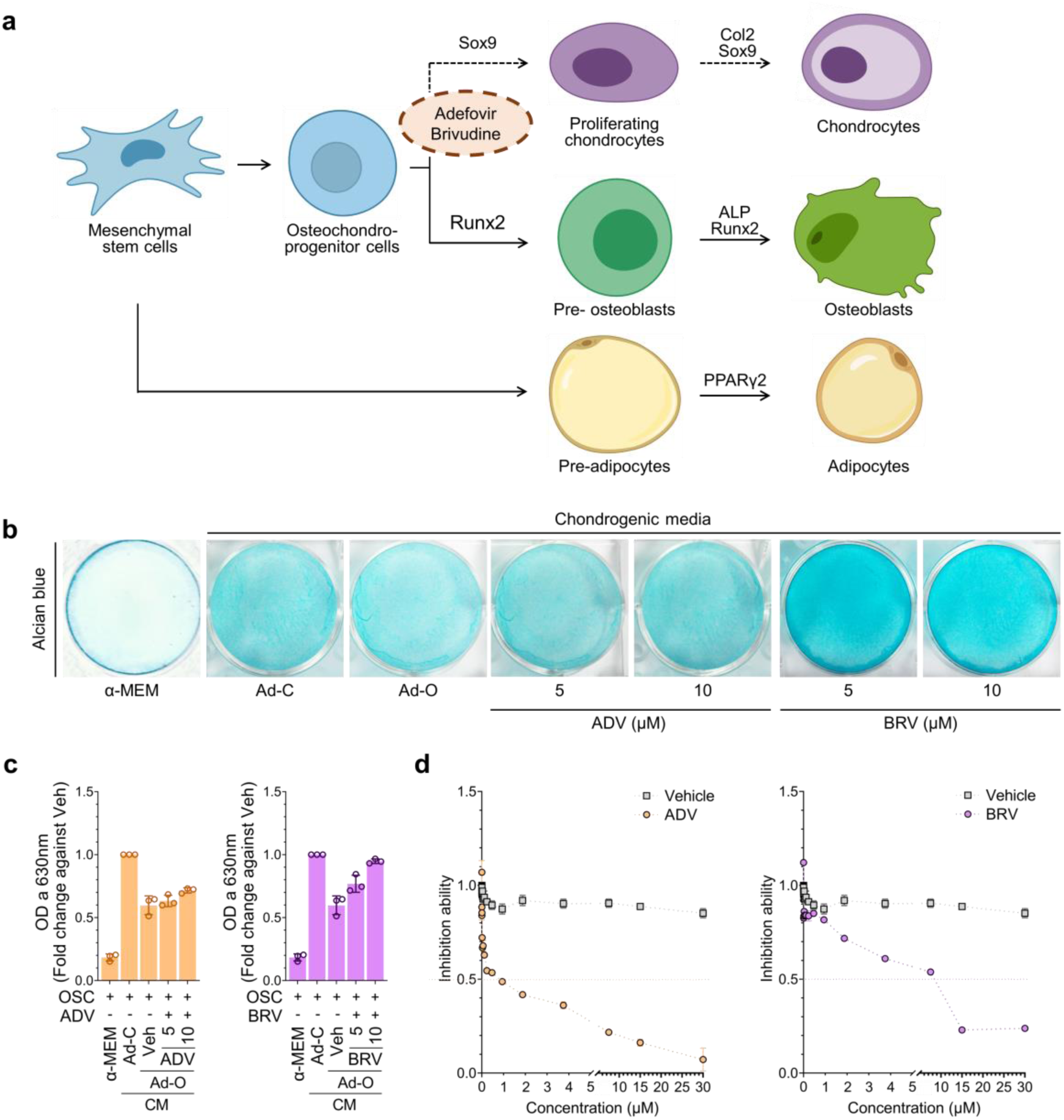
BRV promotes chondrogenic differentiation and supports cartilage regeneration. (a) Schematic of the in vitro assay to assess the chondrogenic potential of ADV and BRV in mesenchymal stem cells (MSCs) isolated from C57BL/6 mouse bone marrow. (b, c) Alcian Blue staining of MSC-derived chondrogenic pellets treated with increasing concentrations of ADV or BRV. BRV induced a stronger and dose-dependent increase in glycosaminoglycan (GAG) production compared to ADV. (d) IC₅₀ values for inhibition of the catabolic marker MMP3 were determined to be 0.48 μM for ADV and 7.5 μM for BRV. Data represent mean ± SEM; *p < 0.05, **p < 0.01 vs. untreated control. The data are shown as means ± s.e.m. *P*-values were obtained by one-way ANOVA followed by Tukey’s multiple comparisons test or two-way ANOVA followed by Sidak’s post hoc test.

### BRV selectively counteracts IL-1β–induced catabolic transcriptional programs in chondrocytes

Treatment with ADV and BRV modulated overlapping but distinct subsets of IL-1β–induced differentially expressed genes (DEGs) in chondrocytes (Figure 7a). At the gene level, IL-1β robustly upregulated matrix-degrading enzymes, including MMP3, MMP9, MMP13, and ADAMTS5, while concomitantly suppressing key anabolic cartilage markers such as ACAN and COL2A1^6,9^ (Figure 7b). BRV consistently reversed these IL-1β–driven transcriptional changes, leading to reduced expression of matrix-degrading enzymes and partial restoration of anabolic cartilage-associated genes. In contrast, ADV showed comparatively limited effects on these gene-level alterations. These BRV-specific transcriptional changes are consistent with a shift away from an extracellular matrix–degrading phenotype characteristic of inflammatory cartilage degeneration.

**Figure 7.**
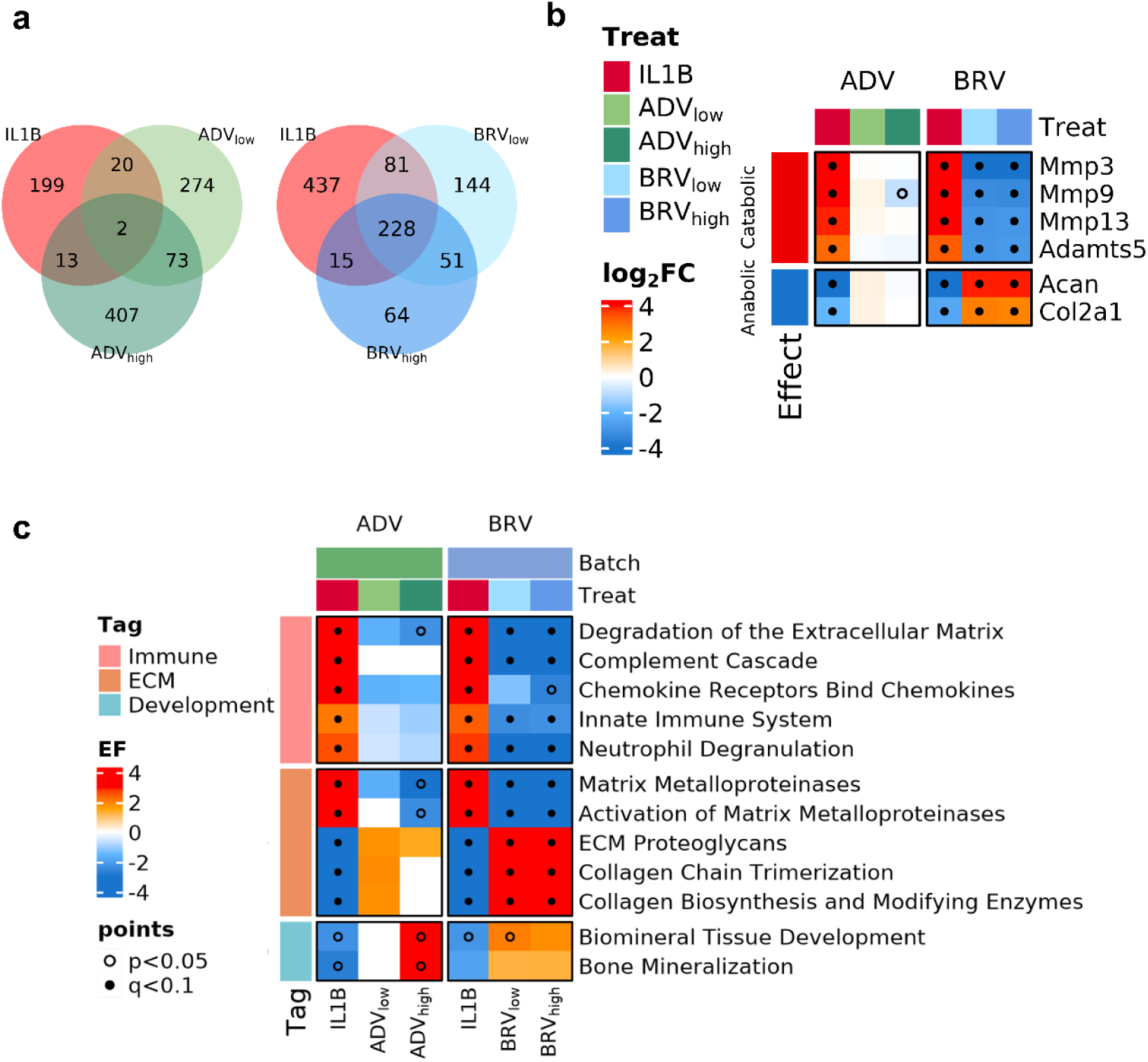
Transcriptomic profiling reveals BRV-mediated attenuation of IL-1β–induced inflammatory and catabolic programs. (a) Venn diagrams illustrating the overlap of differentially expressed genes (DEGs) induced by IL-1β and their modulation by low- and high-dose treatments of ADV(left) or BRV(right). (b) Heatmap showing expression changes of representative catabolic and anabolic cartilage-associated genes across treatment conditions. (c) Pathway enrichment analysis highlighting IL-1β–activated inflammatory and matrix-degrading pathways and their modulation by ADV and BRV. P values were obtained from one-sided hypergeometric tests and adjusted by the Benjamini–Hochberg procedure; dots indicate statistical significance (*p* < 0.05, *q* < 0.1).

At the pathway level, IL-1β exposure induced robust activation of inflammatory and immune-related programs, including complement cascade activation, chemokine receptor signaling, innate immune responses, and neutrophil degranulation (Figure 7c).

In parallel, pathways associated with ECM proteoglycans, collagen biosynthesis and modification, which related with cartilage structural maintenance^5^ were strongly suppressed. Both compounds partially attenuated IL-1β–induced pathway-level alterations, however, BRV exerted more pronounced suppression of inflammatory programs and stronger reactivation of cartilage-associated pathways compared to ADV.

Collectively, these data indicate that BRV effectively counteracts IL-1β–induced catabolic remodeling by suppressing inflammatory and matrix-degrading transcriptional programs while promoting the reactivation of anabolic signaling networks essential for cartilage integrity.

### ADV- and BRV-Associated Changes in Chondrocyte Viability and Transcriptomic Profiles

Treatment of IL-1β–stimulated chondrocytes with ADV and BRV resulted in a dose-dependent increase in cell viability (Figure 8a), with BRV conferring greater protective effects than ADV. To investigate the molecular basis underlying this phenotypic improvement, we examined transcriptomic changes associated with compound treatment under inflammatory conditions.

**Figure 8.**
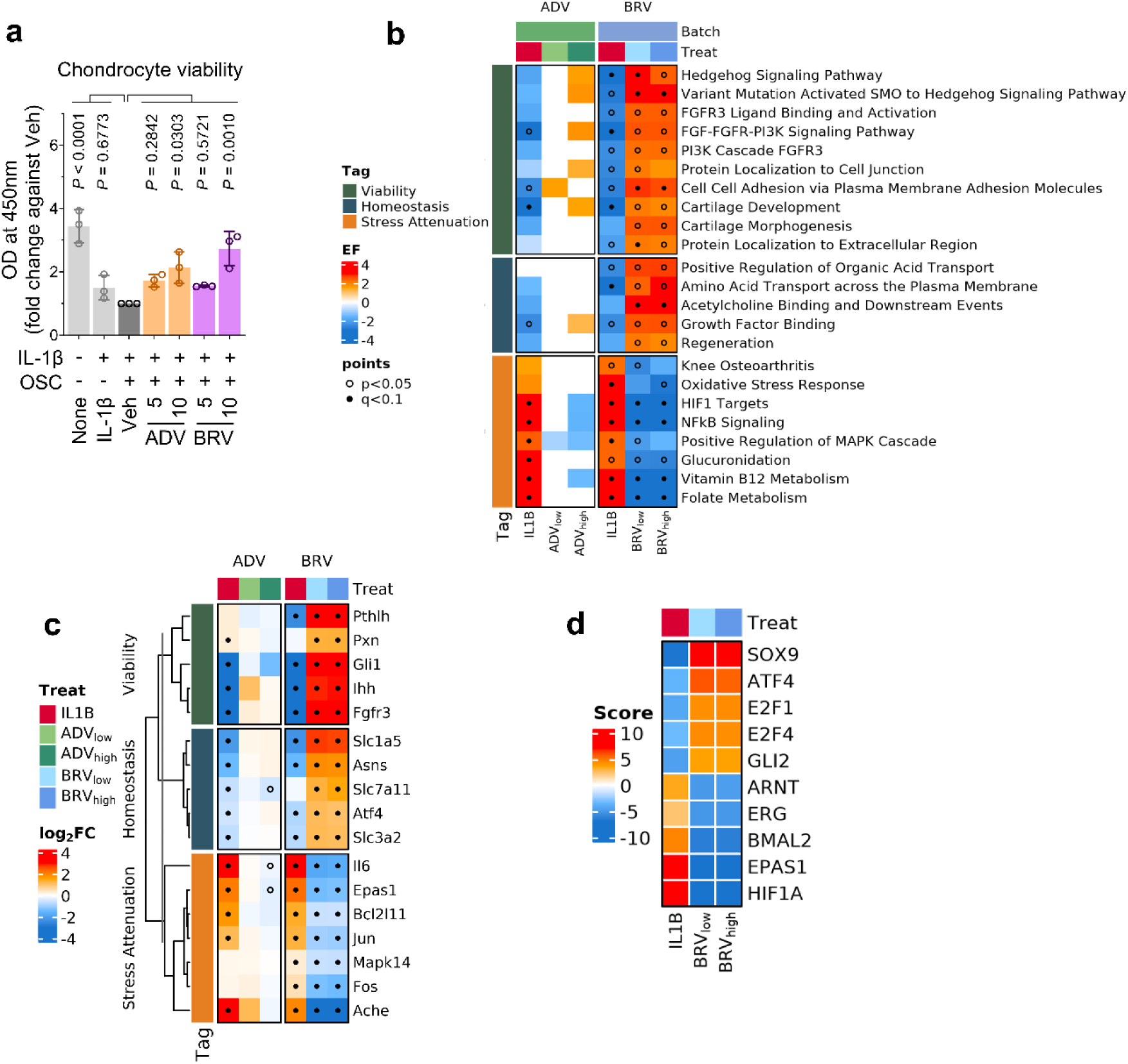
BRV-associated transcriptional programs supporting chondrocyte viability under inflammatory stress. (a) Cell survival assays demonstrating that treatment with ADV and BRV significantly increased the viability of primary chondrocytes. *P*-values were obtained by one-way ANOVA followed by Tukey’s multiple comparisons test or two-way ANOVA followed by Sidak’s post hoc test. (b) Pathway-level enrichment analysis highlighting BRV-associated restoration of developmental and homeostatic pathways and suppression of stress-related signaling programs. P values were calculated using one-sided hypergeometric tests and adjusted by the Benjamini–Hochberg procedure. Dots indicate statistically significant pathways (p < 0.05, q < 0.1). (c) Heatmap showing expression changes of representative genes grouped into viability, homeostasis, and stress-associated categories. (d) Transcription factor activity analysis illustrating BRV-associated regulatory shifts toward chondrogenic identity and cellular maintenance.

Pathway-level analysis revealed that BRV reversed IL-1β–induced suppression of programs associated with chondrocyte viability and functional maintenance^21,22^, including Hedgehog signaling, FGFR3-associated pathways, cartilage development, and cell–cell interaction processes (Figure 8b).

In addition, BRV treatment was associated with upregulation of cellular homeostasis–related pathways, such as amino acid transport^23^ and acetylcholine binding^24^, which support metabolic balance, stress adaptation, and long-term maintenance of chondrocyte function.

In parallel, BRV dampened stress-related signaling pathways, including oxidative stress responses^25^, NF-κB signaling^26^, and MAPK cascade activation^27^. Notably, IL-1β–induced enrichment of stress-adaptive metabolic programs—such as glucuronidation and one-carbon metabolism (folate and vitamin B12 pathways)—was also attenuated by BRV, suggesting normalization of secondary metabolic responses associated with inflammatory stress^28^.

To resolve the gene-level composition underlying the pathway-level alterations described above, we examined representative genes contributing to the enriched viability- and homeostasis-associated pathways. Consistent with the pathway-level signatures, BRV treatment upregulated representative genes associated with chondrocyte viability and identity maintenance, including key regulators of Hedgehog and FGFR3 signaling, as well as genes involved in amino acid transport and metabolic support (Figure 8c). These genes constitute core components of the developmental, metabolic, and homeostatic pathways restored by BRV under inflammatory conditions^21,23,29^.

Conversely, genes contributing to inflammatory burden, apoptotic priming, and degenerative signaling^30–33^—represented by cytokines and stress-responsive signaling components—were consistently suppressed, reflecting attenuation of stress-associated transcriptional programs enriched at the pathway level.

Transcription factor activity analysis inferred by decupler^34^ further supported this coordinated regulatory shift (Figure 8d). BRV treatment was associated with increased inferred activity of transcription factors governing chondrogenic identity and proliferative competence^7,23,35^—including SOX9, ATF4, E2F family members (E2F1 and E2F4), and GLI2—while reducing the activity of stress- and hypoxia-associated regulators such as HIF1A, EPAS1, BMAL2, ERG, and ARNT^36–38^. Notably, these hypoxia-associated factors were elevated under IL-1β stimulation, and their attenuation by BRV occurred alongside suppression of inflammatory and oxidative stress pathways, suggesting a rebalancing of stress-responsive transcriptional programs rather than complete abrogation of hypoxia-related signaling.

Collectively, these results indicate that BRV does not act through a single pathway, but rather rebalances the transcriptional landscape toward enhanced chondrocyte viability and functional homeostasis under inflammatory stress.

## Discussion

Osteoarthritis (OA) remains a prevalent and progressive joint disorder for which no approved disease-modifying therapies are currently available. Although the OSCAR has emerged as a key regulator of cartilage homeostasis and OA pathogenesis, its therapeutic exploitation has been limited by the perceived difficulty of targeting collagen-recognizing receptors with small molecules. In this study, we demonstrate that OSCAR can be effectively modulated using a structure-independent, bioactivity-driven discovery strategy and identify two repurposable compounds, ADV and BRV, that attenuate OA-associated pathological processes across molecular, cellular, and organismal levels.

A central conceptual advance of this work lies in overcoming intrinsic limitations associated with OSCAR as a drug target. Unlike classical enzymes or receptors that possess deep and well-defined ligand-binding pockets, OSCAR recognizes collagen through an extended, surface-exposed interface within its membrane-proximal D2 domain. Such interaction modes are typically refractory to conventional structure-based drug discovery approaches. By leveraging sBEAR, a data-centric framework that mines large-scale bioactivity data rather than relying on explicit structural templates or known ligands, we were able to prioritize candidate compounds predicted to interfere with OSCAR–collagen engagement despite the absence of canonical binding pockets. This highlights the utility of large-scale bioactivity information as a complementary discovery axis for targets that have historically proven difficult to address using target-centric strategies.

The robustness of this discovery strategy is supported by the consistency of experimental validation across multiple biological scales. Candidate compounds prioritized by sBEAR inhibited OSCAR–collagen binding in cell-free assays and attenuated IL-1β–induced catabolic responses in chondrocytes. Among these, ADV and BRV emerged as lead compounds that reproducibly suppressed OSCAR-associated signaling and preserved cartilage integrity in vivo. In a surgically induced DMM mouse model of OA, intra-articular administration of both compounds significantly preserved cartilage integrity and reduced associated pathological features, when therapeutic intervention was initiated one week after disease induction. The alignment between computational prediction, biochemical inhibition, cellular response, and tissue-level protection provides strong support for the predictive value of the integrated screening and validation pipeline.

Structural docking analyses provided a structural framework for the engagement of chemically distinct small molecules with OSCAR. ADV and BRV were predicted to localize to the collagen-recognition surface of the OSCAR D2 domain, overlapping with residues previously implicated in collagen binding. These observations are consistent with structural studies indicating that OSCAR recognizes collagen through an extended surface interface rather than a discrete binding pocket.

Beyond direct receptor engagement, transcriptomic profiling revealed that OSCAR inhibition reshapes disease-relevant gene regulatory programs in chondrocytes. BRV induced broad attenuation of IL-1β–driven inflammation and catabolic process, while partially restoring anabolic, developmental, and homeostatic programs for cartilage maintenance. These effects were reflected across pathways related to extracellular matrix organization, cellular viability, and stress responses, consistent with broader transcriptional changes associated with OA-related processes. Notably, Hedgehog-related pathways were modulated by BRV as well as their key transcription factors. While Hedgehog signaling was reported to promote chondrocyte hypertrophy and osteoarthritis progression^39^, our data reproduced that its biological roles are highly context- and stage-dependent^29^. Up-regulation genes were also highly enriched with the downstream genes of GLI family transcription factors and SOX9, suggesting their coordinated modulation.

In parallel, BRV attenuated IL-1β–induced stress- and hypoxia-associated regulatory programs, accompanied by reduced activity of HIF1A, EPAS1, and other stress-responsive regulators, as well as suppression of oxidative stress, NF-κB, and MAPK signaling pathways. Given the context-dependent roles of hypoxia-inducible factors in cartilage biology^36^, these changes are consistent with normalization of inflammation-associated stress responses.

In contrast, ADV showed more limited transcriptomic reprogramming despite improving chondrocyte viability and cartilage preservation in vivo, suggesting that phenotypic protection may arise through partially transcription-independent mechanisms. This distinction underscores the value of integrating functional, in vivo, and transcriptomic analyses when interpreting therapeutic effects.

Several limitations of this study should be acknowledged. First, while docking analyses support engagement of the OSCAR D2 domain, direct biophysical measurements will be required to quantify binding affinities and kinetics and to confirm competitive inhibition under physiological conditions. Second, transcriptomic analyses were focused primarily on BRV as a representative lead compound, and thus may not fully capture the molecular mechanisms underlying ADV-mediated protection. Third, although the DMM model recapitulates key features of post-traumatic OA, additional models will be necessary to evaluate the generalizability of OSCAR inhibition across diverse OA etiologies and disease contexts.

In summary, this work establishes OSCAR as a viable therapeutic target in OA and demonstrates that structure-independent, bioactivity-driven screening can uncover repurposable small molecules for receptors previously considered intractable to conventional small-molecule discovery. Beyond the identification of ADV and BRV as candidate disease-modifying agents, our findings highlight a generalizable, data-centric discovery paradigm that leverages large-scale bioactivity information to expand therapeutic opportunities across challenging disease contexts.

## Methods

### Bioactivity-driven virtual screening with the sBEAR framework

To identify small-molecule inhibitors of OSCAR, we employed an expanded version of the BEAR algorithm, termed sBEAR. The BEAR framework repurposes large-scale bioassay data by identifying *hit-enriched assays* (HEAs), defined as assays whose hits significantly overlap with a given query ligand set, and integrates enrichment across assays to prioritize candidate compounds^16^.

sBEAR extends this framework by replacing exact hit overlap with structural similarity-based pairwise enrichment, thereby allowing structurally related—but not identical—compounds to contribute to enrichment signals. This modification enables scaffold hopping and expands predictive coverage to targets with limited known ligands.

For a given protein target P:

Q: Query compound set

H: Hit compound set of a given bioassay

H^c^: Non-hit compound set of a given bioassay

The similarity significance S is calculated as the number of compound pairs between two sets that have a Tanimoto coefficients (Tc) above a specified threshold. For example, S_Q×H_ denotes the number of similar pairs between the query set Q and the hit set H.

A contingency table was constructed as follows:

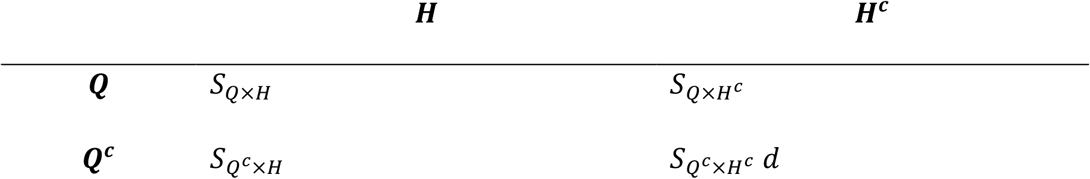

The structurally Similar Hit Enrichment Score (sHES) for each assay was calculated as a log_2_ odds ratio comparing similarity enrichment between 𝑆_𝑄×𝐻_ and 𝑆_𝑄_^𝑐^_×𝐻_ normalized by the background similarity distribution. Assays with sHES greater than a preset cutoff (typically sHES > 1) are designated as HEAs for the query set.

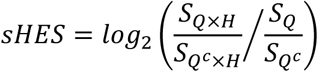

For each HEA, compounds were then ranked by reported bioactivity (e.g. IC_50_, % inhibition) and partitioned into equal-sized activity bins. Bin-specific sHES values are recalculated, and a monotonic regression was applied to enforce consistency between compound rank and enrichment while reducing noise. The regressed sHES were then assigned back to individual compounds. Finally, for every compound appearing in any HEA, each compound’s sBEAR score was defined as the sum of its sHES values across all HEAs in which it was tested.

To screen OSCAR-modulating compounds, known OSCAR ligands from human and six mammalian species (17 and 280 ligands, respectively) were used as the query set. sBEAR was applied to the PubChem BioAssay database comprising 163,220 assays and 1,231,461 compounds^40^. Bioassays enriched for structurally similar ligands were selected as HEAs, and compounds were ranked by their aggregated sBEAR scores to generate the candidate list used for subsequent docking and experimental validation.

### Molecular docking and pose refinement of OSCAR–ligand complexes

The 3-dimensional structure of OSCAR was obtained from the Protein Data Bank (PDB ID: 5CJB)^12^. Ligand structures, including FDA-approved drugs and top-ranked compounds identified from sBEAR screening, were prepared from their SMILES representations. Protonation states were assigned corresponding to physiological pH 7.0, hydrogen atoms were added, and ligand structures were converted to PDBQT format using OpenBabel^41^.

Initial docking-based affinity estimation was performed as described above using AutoDock Vina^19^ to evaluate relative binding propensities across candidate compounds. To further refine binding conformations and explore plausible interaction geometries within the collagen-recognition surface of OSCAR, pose refinement was conducted using Protenix, an AlphaFold3-based structure prediction platform^20^. Protenix simulations were performed by providing the OSCAR amino acid sequence together with the CCD identifiers of selected ligands, enabling flexible modeling of ligand engagement with extended and surface-exposed binding regions that are not well captured by rigid docking approaches.

The resulting 3-dimensional docking poses were subsequently analyzed using ProteinsPlus^42^ to generate two-dimensional interaction maps and to identify key molecular contacts, including hydrogen bonding, electrostatic interactions, and aromatic stacking features. These analyses were used to compare interaction patterns across candidate ligands and to assess their spatial overlap with the previously defined collagen-recognition region within the OSCAR D2 domain.

### Mice

Murine experiments were conducted with male 10–11-week-old C57BL/6J (C57BL/6BomTac, DBL, South Korea) or Institute of Cancer Research (ICR) mice (IcrTac; ICR, DBL, South Korea). All mice were housed in pathogen-free barrier facilities at 5 or less per cage at 24–26°C, humidity ranging at 30–60%, and with a 12 hour light/dark cycle. The mice were randomly allocated to each experimental group. All animal experiments were approved by the Institutional Animal Care and Use Committees (IACUC, Protocol No: IACUC 22-003, 25-006) of Ewha Womans University and followed National Research Council Guidelines.

### ELISA assay for OSCAR binding activity

Collagen was coated onto cell culture plates, and ELISA assays were performed in the presence of hOSCAR-Fc and a compound. For verification, the COL^pep^/hOSCAR-Fc-based ELISA described above was used with increasing concentrations of candidate molecules. Moreover, to determine whether the candidates could compete with collagen-II for binding to cell-surface OSCAR, murine chondrocytes were cultured for 48 h in plates on which 2 μg/ml COL^pep^ had been immobilized, infected with Ad-OSCAR for 2 h, and then treated with each candidate molecule for 48 h. Alternatively, murine chondrocytes were not infected with Ad-OSCAR. OSCAR mRNA/protein was isolated and measured by qRT-PCR.

### Molecule information of materials

ADV (Adefovir) was obtained from Sigma-Aldrich (Cat# SML0240, Sigma-Aldrich, St Louis, MO, USA) and dissolved in PBS. BRV (Brivudine) was obtained from Sigma-Aldrich (Cat# B9647, Sigma-Aldrich, St Louis, MO, USA) and dissolved in PBS. Vector Biolabs (Malvern, PA, USA) manufactured the adenoviruses that expressed OSCAR (Ad-OSCAR; catalog No. ADV-267721), Ad-Control (1060)

### Primary culture of articular chondrocytes and cell line culture

Articular chondrocytes of mice were isolated from femoral condyles and tibial plateaus of 4- to 5-day-old Institute of Cancer Research (ICR) mice by digestion with 0.2 % collagenase type Ⅱ. Culture dishes were pre-coated with OSC collagen peptide (2 μg/ml) or pre-coated with GPP10 control peptide at 4 °C overnight. The chondrocytes were maintained in Dulbecco’s modified Eagle’s medium (DMEM; HyClone, Logan, UT, USA) containing 10 % fetal bovine serum (FBS) and were treated with 10 ng/ml of IL-1β or 5 - 20 μM of ADV and BRV for 48 h. Mouse articular chondrocytes were cultured for 48 hours, infected with the indicated multiplicity of infections (MOIs) of adenovirus for 2 h, and cultured in the presence or absence of pharmacological agents, ADV and BRV for an additional 24 h. Articular chondrocytes were isolated from femoral condyles and tibial plateaus of 2- to 3-day-old WT and Oscar-/- mice. Chondrocytes were seeded at the pre-incubated dish with OSC collagen peptide (2 μg/ml) for 16 h, before being treated with IL-1β or 0.1 μg/ml hOSCAR-Fc fusion protein. Articular chondrocyte apoptosis was determined using the TUNEL assay and a kit from Millipore (Apoptosis Detection kit, Lot #2397039, Temecula, CA, USA). The OSCAR-binding triple-helical peptide was purchased from University of Cambridge (CB2 1QW, UK), and the purity was analyzed using high-performance liquid chromatography. A triple-helical-forming peptide composed of the minimal binding motif GPC-(GPP)5-GPOGPAGFO-(GPP)5-GPC-amide was used for western blotting and quantitative reverse transcription polymerase chain reaction analyses and best dissolved in Acetic acid, 0.01 M^11^. GPP10 collagen-mimetic peptide composed of GPC-(GPP)10-GPC-amide was used as negative control peptide.

### Histological analysis of OA

The knee joints of mice were fixed in 10 % formaldehyde at 4 °C for >24 h, decalcified in 0.5 M ethylenediaminetetraacetic acid in PBS (pH 7.4) for 2 weeks, embedded in paraffin, sliced into 5-μm sections, and stained with hematoxylin and eosin, 0.1% Safranin-O (s8884; Sigma-Aldrich, St Louis, MO, USA), and 0.05% Fast green FCF (f7258; Sigma-Aldrich, St Louis, MO, USA). Sclerosis and articular cartilage destruction were identified by safranin-O staining and measured with OsteoMeasureXP (OsteoMetrics, Inc., Atlanta, GA, USA), Image-pro plus (v4.5, Media Cybernetics, Inc., Rockville, USA), Adobe photoshop (v9.0, San Jose, CA, USA), and an Olympus DP72 charge-coupled device camera (v2.1, Olympus Corporation, Tokyo, Japan). Articular cartilage destruction was scored by using OARSI grades (0–6), which is a standard OA-grading system^43,44^ (Supplementary Table 2). The OARSI grades of the medial tibia were assessed by averaging the scores of three experienced investigators. The ratio of hyaline cartilage to calcified cartilage, osteophyte maturity, and synovitis^45^ were also determined (Supplementary Table 3). Moreover, subchondral bone sclerosis was determined by measuring the SBP thickness.

### RNA isolation and qRT-PCR

Total RNAs from primary chondrocytes or bone marrow-derived MSCs were isolated by using TRIzol (Invitrogen, Carlsbad, CA, USA). Total RNAs from murine and human knee joint tissues were isolated by using the RNA Mini Kit (Life Technologies, Carlsbad, CA, USA). The RNAs were reverse-transcribed to generate complementary DNA (cDNA) by using the Superscript cDNA synthesis kit (Invitrogen) according to the manufacturer’s instructions. qRT-PCR was performed using the KAPA SYBR Green fast qPCR kit (Kapa Biosystems, Inc., Wilmington, MA, USA) on a Step One Plus RT-PCR machine (Applied Biosystems, Foster City, CA, USA). The samples were tested in triplicate and the data were normalized by using β-actin as a housekeeping gene. Supplementary Table 4 provides the full list of primers.

### Experimental OA in mice

C57BL/6J (WT) male mice were used for experimental OA studies. We purchased 10-to 11-week-old C57BL/6 male mice from Japan SLC, Inc. (Hamamatsu, Japan); Experimental OA was induced by DMM (Destabilized of medial meniscus) surgery in 10-to 11-week-old male mice. In DMM surgery, the medial meniscus ligament was surgically removed from the right knee joint of hind limb. Mice injected with vehicle or sham-operated mice were used as controls. A sham operation was performed on the contralateral knee with no DMM surgery. To study the protective effects of ADV and BRV from induced OA, ADV and BRV were dissolved in DMSO. Mice were given an IA (Intra-articular) injection of 0.5 and 2 mg/kg of ADV and BRV or 2 mg/kg of a vehicle control once per week for 8 weeks, beginning one week after DMM surgery, respectively. The mice were euthanized at 9 weeks after OA induction and subjected to biochemical and histological analyses. All mice were maintained in pathogen-free barrier facilities. Mice were housed in barrier facility at 5 or less per cage at 24 - 26 °C with humidity ranging between 30 - 60 % with 12 hour light/dark cycle. For each experiment, age-and sex-matched mice were used and randomly allocated to each experimental group. All animal experiments were approved by the Institutional Animal Care and Use Committees (IACUC Protocol No: IACUC 22-003, 25-006) of Ewha Womans University and followed National Research Council Guidelines.

### Generation of bone marrow-derived MSCs and chondrogenesis

Primary bone marrow mesenchymal stem cells were isolated from the tibias and femurs of 4–5-week-old male C57BL/6J mice as previously described^46^. The cells were then incubated for 3 weeks in chondrogenic medium (Hyclone, α-MEM containing 10 µg/ml insulin, 10 µg/ml transferrin, and 6 µg/ml sodium selenite) with or without ADV and BRV (10 µM).

### Alcian-Blue staining of cartilage glycosaminoglycans and chondrogenic MSCs

Bone marrow-derived MSCs cultured in chondrogenic medium were stained with Alcian Blue (B8438; Sigma Aldrich) and photographed with Olympus DP72 charge-coupled device camera (v2.1, Olympus Corporation, Tokyo, Japan). The Alcian-Blue activity in the dishes was also quantified by measuring the absorbance at 630 nm.

### RNA sequencing analysis

Primary chondrocytes were cultured for 2 days and infected with Ad-OSCAR at a MOI of 800 for 2 h. Following infection, cells were cultured for an additional 24 h in the presence or absence of ADV or BRV (10 µM). Control cells were infected with Ad-Control under identical conditions. All experiments were performed on culture plates coated with COL^pep^. Total RNA was extracted from chondrocytes, and sequencing libraries were prepared using the TruSeq Stranded mRNA Library Prep Kit (Illumina, San Diego, CA, USA) according to the manufacturer’s instructions. Paired-end RNA sequencing (2 × 100 bp) was performed on an Illumina NovaSeq 6000 platform. Raw sequencing quality was assessed using FastQC, and adapter trimming and quality filtering were performed with BBDuk. Processed reads were aligned to the mouse reference genome (GRCm38) using STAR (v2.7.9a), and gene-level raw read counts were quantified with RSEM (v1.3.1).

Raw and processed RNA-seq data have been deposited in the Gene Expression Omnibus (GEO) under accession number GSE298720. Differential gene expression analyses were conducted using DESeq2. Comparisons were performed between control chondrocytes and IL-1β–treated chondrocytes, as well as between IL-1β–treated chondrocytes and those treated with ADV or BRV. Multiple testing correction was applied using the Benjamini–Hochberg procedure^47^.

Given the distinct magnitude and distribution of transcriptional responses induced by IL-1β, BRV, and ADV, different cutoff criteria were applied to define differentially expressed genes (DEGs). For IL-1β and BRV, DEGs were defined as genes with |log₂ fold change| > 3 and false discovery rate (FDR) < 0.1. For ADV, which exhibited more modest transcriptional effects, DEGs were defined using |fold change| > 1.5 and nominal p-value < 0.05.

TF activity was inferred using the weighted mean method implemented in decoupler (v.1.6.0)^34^, based on TF–target interaction networks obtained from DoRothEA using confidence levels A–C.

### Statistical analysis

The sample size for each experiment was not predetermined. To analyze differences between 2 and >2 groups, Student’s two-tailed *t*-test, or one-way analysis of variance (ANOVA), or two-way ANOVA were conducted, respectively. If ANOVAs were significant, pairwise multiple comparisons were conducted. Data based on the comparison of two samples with a variable measured using an ordinal grading system were analyzed using the Sidak’s multiple comparisons test. Data based on ordinal grading systems were analyzed using the non-parametric Mann–Whitney *U* test. Each n indicates the number of biologically independent samples, mice per group or human specimens. The sample size for each experiment was not predetermined. The *P* values are indicated in the figures or in Source Data, and the error bars represent s.e.m. for parametric data and the calculated 95% CIs for nonparametric data. Except where stated, the experiments were not randomized and the investigators were not blinded to allocation during experiments and outcome assessment. Multiple comparisons were performed using Tukey’s test, or Sidak’s test, or Dunn’s test with *P* values set at <0.0001. All graphs and statistical analyses were made by using GraphPad Prism (v8.4.3, San Diego, CA, USA). *P-*values are indicated in the figures. The error bars in the figures show s.e.m. or 95% confidence intervals. All data were collected from at least three independent experiments.

## Competing interests

The authors declare that they have no competing interests.

## Data and materials availability

RNA-seq data of the IL-1β and ADV, BRV treated chondrocytes have been deposited in the GEO under accession number GSE298720.

## Acknowledgements

This research was supported by the National Research Foundation of Korea (NRF) grant funded by the Basic Science Research Program through the National Research Foundation of Korea by the Ministry of Science and ICT (NRF-2022R1A2C1091260, NRF-RS-2025-00519443, NRF-RS-2025-25427404, NRF-RS-2023-00217798).

## Authors contributions

WK and SYL conceived, designed, and supervised the study. GR, SP and WK performed the computational analyses, data interpretation, and in silico predictions. JK conducted the wet-lab experiments. WK and GR integrated the results and drafted the manuscript. All authors contributed to manuscript revision and approved the final version.

## Notes

### Competing Interest Statement

The authors have declared no competing interest.

