## Supplementary Information for "Bioactivity-driven discovery of repurposable antivirals as OSCAR inhibitors that promote cartilage protection via transcriptomic reprogramming"

### Bioactivity-driven discovery identifies repurposable OSCAR inhibitors for disease modification in osteoarthritis

* Corresponding Author

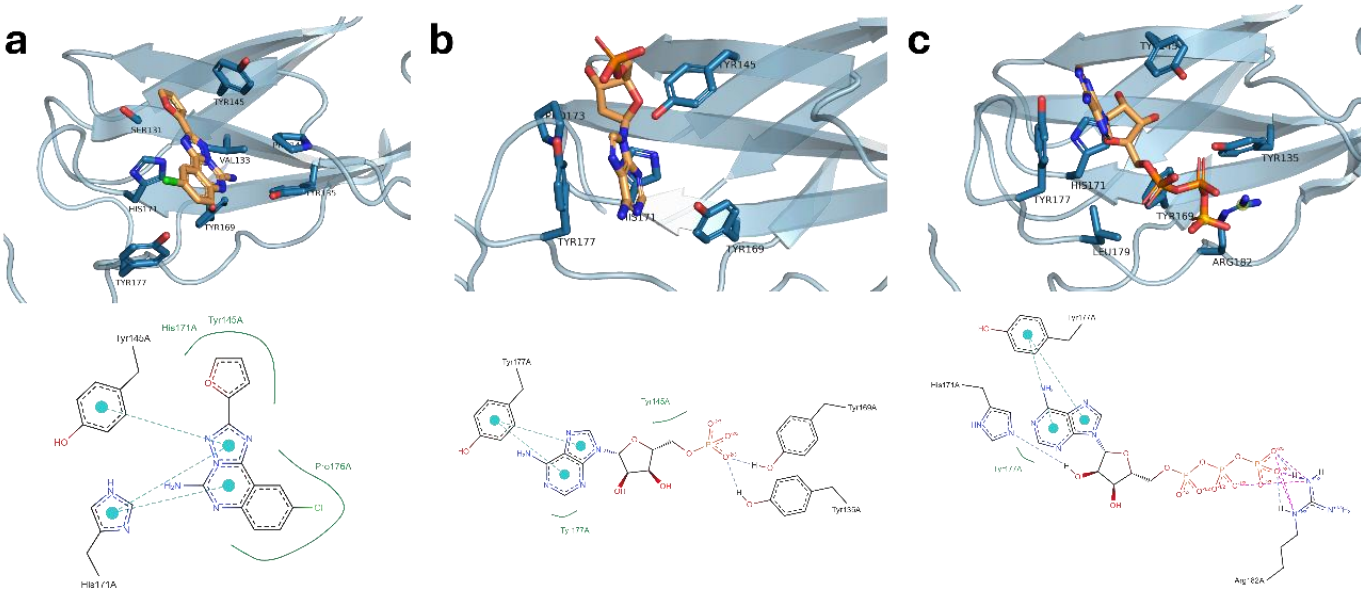


**Supplementary Figure 1. Consistent docking of additional top-ranked compounds to the OSCAR D2 collagen-recognition surface.**

Predicted binding poses (top) and corresponding two-dimensional interaction maps (bottom) of CGS-159 (a), AMP (b), and ATP (c) docked to the OSCAR D2 domain. All compounds localize the OSCAR D2 collagen-recognition surface, consistent with engagement of a shared interaction region across structurally diverse candidates. These compounds represent chemically distinct nucleotide-like or purine-based structures identified among the top-ranked candidates.

**Supplementary Table 3. The recommended semi-quantitative scoring system.**

| Grade | Osteoarthritic Damage |
|---|---|
| 0 | Normal |
| 0.5 | Loss of Safranin-O without structural changes |
| 1 | Small fibrillations without loss of cartilage |
| 2 | Vertical clefts down to the layer immediately below the superficial layer and some loss of surface lamina |
| 3 | Vertical clefts/erosion to the calcified cartilage extending to <25% of the articular surface |
| 4 | Vertical clefts/erosion to the calcified cartilage extending to 25-50% of the articular surface |
| 5 | Vertical clefts/erosion to the calcified cartilage extending to 50-75% of the articular surface |
| 6 | Vertical clefts/erosion to the calcified cartilage extending to >75% of the articular surface |

**Supplementary Table 4. Criteria for measuring and quantifying synovitis.**

| Grade | Enlargement of the synovial lining cell layer |
|---|---|
| 0 | The lining cells form one layer |
| 1 | The lining cells form 2-3 layers |
| 2 | The lining cells form 4-5 layers, few multinucleated cells might occur |
| 3 | The lining cells form more than 5 layers, the lining might be ulcerated and multinucleated cells might occur |

| Grade | Density of the resident cells |
|---|---|
| 0 | The synovial stroma shows normal cellularity |
| 1 | The cellularity is slightly increased |
| 2 | The cellularity is moderately increased, multinucleated cells might occur |
| 3 | The cellularity is greatly increased, multinucleated giant cells, pannus formation and rheumatoid granulomas might occur |

| Grade | Inflammatory infiltrate |
|---|---|
| 0 | No inflammatory infiltrate |
| 1 | Few mostly perivascular situated lymphocytes or plasma cells |
| 2 | Numerous lymphocytes or plasma cells, sometimes forming follicle-like aggregates |
| 3 | Dense band-like inflammatory infiltrate or numerous large follicle-like aggregates |

| Sum | Summer grade |
|---|---|
| 0 or 1 | No synovitis |
| 2 to 4 | Low-grade synovitis |
| 5 or 6 | Middle-grade synovitis |
| 7 to 9 | High-grade synovitis |

**Supplementary Table 5. Oligonucleotides used for RT-PCR and real-time quantitative PCR analysis.**

| Primer | | | | Sequence (5' to 3') | |
|---|---|---|---|---|---|
| **RT-PCR** | OSCAR WT | F | 5' | GCTTCTTCTTTGCAGCTCCT | 3' |
| | | R | 5' | ATCGTCATCCATGGCGA | 3' |
| | OSCAR KO | F | 5' | CAAAGCCAGAGTCCTTCAGA | 3' |
| | | R | 5' | GATGGTCTTGGTCCTTAGCC | 3' |
| **real-time quantitative PCR** | OSCAR | F | 5' | GCTTCTTCTTTGCAGCTCCT | 3' |
| | | R | 5' | ATCGTCATCCATGGCGA | 3' |
| | MMP3 | F | 5' | TCCTGATGTTGGTGGCTTCAG | 3' |
| | | R | 5' | TGTCTTGGCAAATCCGGTGTA | 3' |
| | MMP13 | F | 5' | TGATGGACCTTCTGGTCTTCTGG | 3' |
| | | R | 5' | CATCCACATGGTTGGGAAGTTCT | 3' |
| | COL2A1 | F | 5' | CACACTGGTAAGTGGGGCAAGA | 3' |
| | | R | 5' | GGATTGTGTTGTTTCAGGGTTCG | 3' |
| | Aggrecan | F | 5' | CTGTCTTTGTCACCCACACAT | 3' |
| | | R | 5' | GAAGACGACATCACCATCCAG | 3' |
| | SOX9 | F | 5' | CACTGGCAGTTACGGCATCAG | 3' |
| | | R | 5' | CATGTAAGTGAAGGTGGAGTAGAGC | 3' |
| | β-Actin | F | 5' | ATATCGCTGCGCTGGTCGTC | 3' |
| | | R | 5' | AGGATGGCGTGAGGGAGAGC | 3' |
